# Expanding the coverage of spatial proteomics

**DOI:** 10.1101/2023.01.29.526114

**Authors:** Huangqingbo Sun, Jiayi Li, Robert F Murphy

## Abstract

Multiplexed protein imaging methods provide valuable information about complex tissue structure and cellular heterogeneity. However, the number of markers that can be measured in the same tissue sample is currently limited. In this paper, we present an efficient method to choose a minimal predictive subset of markers that for the first time allows the prediction of full images for a much larger set of markers. We demonstrate that our approach also outperforms previous methods for predicting cell-level marker composition. Most importantly, we demonstrate that our approach can be used to select a marker set that enables prediction of a much larger set that could not be measured concurrently.

## Main

Multiplexed imaging methods enable researchers to analyze individual cell properties and their spatial relationships in complex tissues. These methods, like co-detection by indexing (CODEX), allow imaging of many dozens of markers in the same tissue sample (1). However, unlike spatial transcriptomics, in which many thousands of different gene transcripts can be measured, spatial proteomics allows only a small fraction of all proteins to be imaged in the same sample.

One approach for addressing this problem is ‘*in silico* labeling’, in which deep learning models are used to predict unmeasured signals from easily acquired reference signals. Examples include predicting subcellular components from unlabeled microscope images (2–5), virtual histological staining of tissue images (6–8), and predicting immunofluorescence or directly inferring cell types from immunohistochemically stained images (9, 10). This concept can be extended to predict a large number of biomarkers from images of a smaller number (11, 12). For example, Wu et al. (13) described a method to select 7 markers out of 40 that enabled accurate prediction of cell types in a number of tissues, and showed the effectiveness of the approach by imaging only those 7.

In this work, we first sought to develop a flexible approach for finding a small subset of markers and using them to predict the full-image expression pattern of the remaining markers. In contrast to the method of Wu et al. (13), we consider the problem of marker selection from an optimization standpoint; our single-panel setting focuses on selecting biomarkers by explicitly modeling the spatial relationship between marker intensities with a graphical model and using a neural network to directly infer the multiplexed fluorescence image instead of predicting expression at the single-cell level. Our basic approach is illustrated in Figure 1a.

**Fig. 1.**
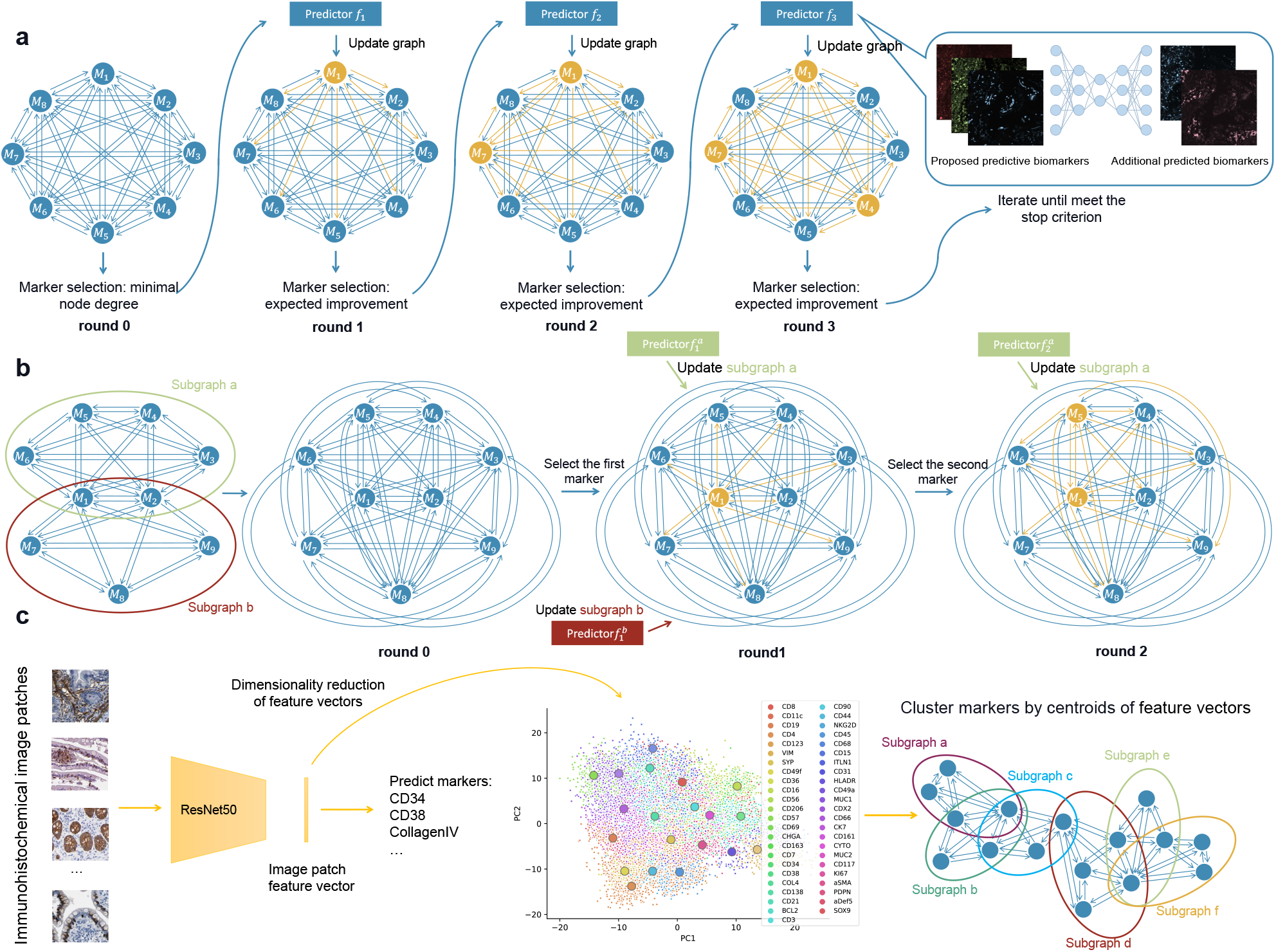
Overview of our methods for identifying predictive markers. **a** Single-panel setting. A clique is constructed with a node for each marker. Then, nodes are iteratively added to a predictive subset (shown in yellow) based on their expected improvement (see Methods), a predictor is trained from that subset (illustrated on the right), and edges attached to them are updated by the predictor’s performance (also shown in yellow). **b** Multipanel setting. The set of markers is partitioned into panels (two in this case) with some overlap ({*M*_1_, *M*_2_ }). Predictors are then constructed from each panel separately. In Round 0, the graph is completed by assigning upperbound edge loadings to edges connecting markers in different panels using the triangle inequality. Markers are iteratively chosen using the graph to be added to the predictive set and predictors are retrained for one or both panels as appropriate. **c** Panel creation. Markers are partitioned into smaller panels based on the centroids of their associated feature vectors extracted with a trained immunohistochemical image classifier.

To evaluate our approach, we used spleen and lymph node CODEX image datasets from the HuBMAP project (14). As described in Methods, each image was first normalized and randomly cropped into small patches. Given a subset of markers, predictors were trained by minimizing the mean square error (MSE) between the predicted and observed marker intensities (see Methods). Visual examination of examples of real and predicted image patches (Figure 2a) for the markers that are predicted the best reveals their patterns are essentially indistinguishable, and that those with the worst predictions are still quite similar. We numerically compared predictions using our selected sets with those of the sets described by Wu et al. (13) and found that our method yields smaller differences between predicted and real images for both tissues (Figure 2b). (For context, since channel intensities were normalized to z-scores, the expected MSE for random predictions within a range of 2 standard deviations is approximately 1.15; since it is calculated over thousands of pixels the chance of obtaining MSE values below 0.3 at random is infinitesimal.) We observed that the errors for the lymph node images were larger than for spleen when using only 7 input channels (the number chosen by Wu et al. (13)). (Principal component analysis (PCA) (S Figure 2) showed that the lymph node profile has a longer tail in its channel eigen-spectrum indicating greater interchannel variability). We therefore selected 5 additional predictive markers for that tissue; the improvement at each step in the training processes is shown in Figure 2c and leads to a better overall MSE. We also characterized the predictions for individual cells (via difference in average channel intensities or in correlations between intensities of pairs of channels) (Figure 2d). Overall, our method achieves better performance in reconstructing single-cell profiles derived from predicted images.

**Fig. 2.**
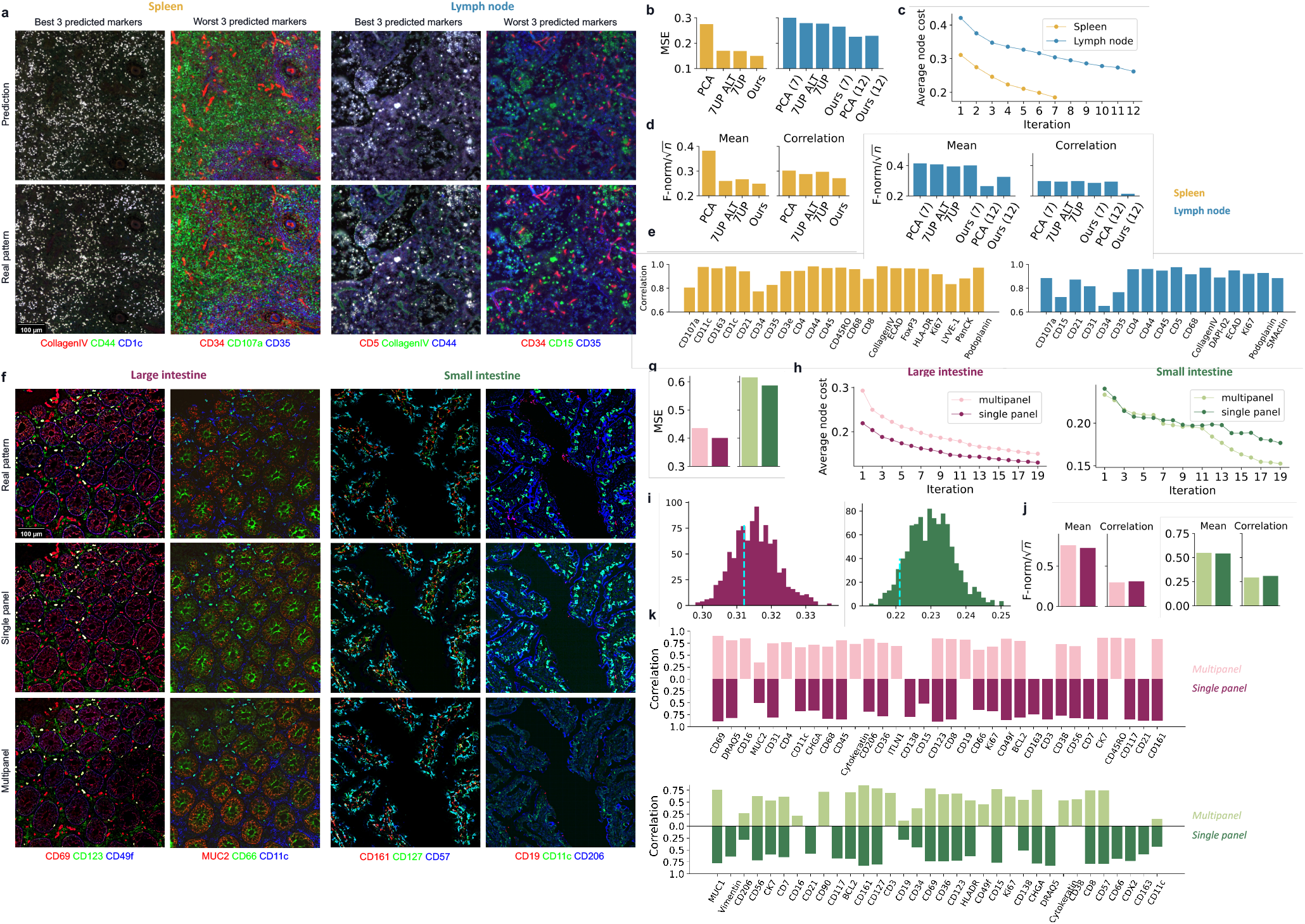
Comparison of marker predictions. Example patches from synthetic and real test images for the best and worst 3 predicted protein markers for spleen and lymph node (**a**), or large and small intestine (**f**). **b, g**, the changes of the average node cost (the overall unpredictability to be minimized) throughout the marker selection procedure. **c, h** reconstruction error (MSE) for various prediction approaches. PCA refers to simply selecting a predictive set using the markers with the most variance in intensity. For lymph node, the number inside the parentheses indicates the number of markers selected. **d, j**, single-cell level assessment (lower values are better). **e, k**, the Pearson Correlation Coefficient (PCC) between synthetic and real images for each marker. **i**, Distribution of average dissimilarity between expression patterns of pairs of markers within each randomly partitioned panel (lower is better). We also mark the resulting average dissimilarity within panels partitioned using immunohistochemical images (cyan dashline). For panel **f**, the worst 3 markers were chosen from the intersection of predicted markers for the single and multi-panel settings.

As discussed earlier, the number of protein markers that can currently be imaged on the same sample is in the dozens, while mammalian cells express tens of thousands of proteins. Thus, creating a predictive set cannot be done by first collecting an image of all (or even hundreds of) markers. We therefore explored a more complicated setting for choosing a predictive set using images of different samples of the same tissue labeled with different panels of markers (see Figure 1b). The first step in this multipanel setting is to choose how to divide a desired large set of markers into smaller panels. We did this using information extracted from independently acquired immunohistochemical images (from Human Protein Atlas (HPA) (15)) (Figure 1c). Markers were partitioned into 5 panels by similarities in their expression patterns (with some overlap; details of panel design are reported in Supplementary Files). Figure 2i shows that the average dissimilarity between markers within individual panels partitioned using immunohistochemical similarities is lower than the average achieved by random partitioning (and quite a bit lower for small intestine). Given just the channels in each panel, we selected a subset of markers by estimating how well that set would be able to predict all markers. We tested this approach using small and large intestine CODEX datasets with 46 markers and assumed that we could only image 19 markers per sample. For each dataset, we first made a holdout test set, then the remaining images were partitioned into training and validation sets (see Methods). To benchmark performance, we ran the single-panel method on all 46 markers as an empirical upper bound for the multipanel method. The difference in test MSE between the two settings is small (Figure 2g). We note that prediction tasks on both intestine datasets are harder than the spleen and lymph node datasets since more markers are required to explain the same fraction of variance (Figure S2).

In Supplementary Files, we listed the markers selected by either the single-or multi-panel approach. For both datasets, a large proportion of all markers are selected in both cases (12 and 13 out of 19). As shown in Figure 2h, though the starting points of the optimization objective are higher for the multipanel setting, it drops faster and goes even lower compared to the single-panel setting in the final iterations. The performances of two settings are very close, and the multipanel was even better for small intestine under some metrics. When example real and predicted patches are examined visually (Figure 2f), both single and multipanel predictors do well even for the worst predicted markers (with the exception of the multipanel predictions for small intestine; the intensities are much lower, but the spatial distributions are still correctly predicted). The very similar performance between single and multipanel is also seen at the single-cell level (Figure 2j).

Lastly, we trained new predictors on datasets from a different batch of experiments using our selected markers to see the generalization performance of our marker choices. We used the 19 markers to train the new predictors on the latest published CODEX image sets of large and small intestine. Note that the new images have 54 markers, more than the number used in the marker selection. Two new predictors were trained using the new datasets, and their generalization performances quantified by the test MSE were 0.591 and 0.536 (single-panel and multipanel) for the new large intestine dataset and 0.563 and 0.583 (single-panel and multipanel) for the new small intestine dataset respectively (roughly similar to our previous results even though the new datasets included 8 previously unseen markers). These results indicate good generalization of the selected sets to new images.

Our results strongly suggest the feasibility of constructing spatial networks for all proteins without imaging them in the same sample and synthesizing multiplexed protein images with high quality. The resulting models would be expected to shed light on protein and cell type spatial interactions in complex tissues. A future direction for optimizing marker panels is to integrate other types of proteomics or genomics information as a prior for panel partitioning.

## Availability

All code and intermediate results are available in a Reproducible Research Archive at https://github.com/murphygroup/CODEXPanelOptimization; it includes a script to download relevant CODEX image data from https://portal.hubmapconsortium.org/.

## Acknowledgements

We thank Tianyue Zhang and Xuecong Fu for helpful discussions during this project. This work was supported in part by grants from the National Institutes of Health Common Fund, OT2 OD026682 and OT2 OD033761.

## Methods

### Data-driven protein marker panel design

First, we introduce the problem and an overview of our proposed heuristic approach. Formally, we denote biomarkers of interest as a set of random variables (RVs) 𝒳 = {*X*_1_, …, *X*_*N*_}, and our goal is to partition 𝒳 into 𝒳_obs_ and 𝒳_pred_ ⊆ 𝒳\ 𝒳_obs_, where 𝒳_obs_ denotes the set of predictive markers and 𝒳_pred_ denotes the set of markers to be predicted. We write the partition as *δ* : 𝒳 →{0, 1}, let *δ*_*i*_ denote *δ*(*X*_*i*_). That is, 𝒳_obs_ = {*X*_*i*_ : *δ*_*i*_ = 1}. Given a partition, the next step is to construct a predictor *f* ∈ ℱ: 𝒳_obs_ → 𝒳_pred_. In this work, we use empirical risk minimization (ERM) to optimize *f*. We denote →_obs_ as the distributions over 𝒳_obs_. With the choice of cost function used in the learning algorithm, *L*, we write the risk of our predictor as ℛ= 𝔼 _(*x,y*)_ ∼ 𝒟 _obs_ × 𝒟_pred_ *L*(*f*). Then, the problem of finding a minimal predictive set can be viewed as the following optimization programming,

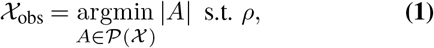

where *ρ* denotes the chosen stopping criterion and 𝒫denotes the powerset.

In general, finding an exactly minimal set of markers is computationally hard.

### Single panel

Given the hardness of the problem, we used a heuristic algorithm. Our algorithm starts with constructing a directed graph 𝒢 = (𝒱, *ε*) where nodes in 𝒱 are associated with the RVs in 𝒳. We use nodes *v*_*i*_ ∈ 𝒱 and RV *X*_*i*_ inter-changeably in the following text. *E* denotes the edge set of 𝒱 where ε = {*e*_*i,j*_ = (*X*_*i*_, *X*_*j*_) : ∀*i, j* ∈ [| 𝒳 |], *i* ≠ *j*}, where each edge has a non-negative *loading w*_*i,j*_. Assume we have some dissimilarity measure *ξ* over 𝒳 × 𝒳, we then initialize the edge loading by the dissimilarity between the RVs associated with the nodes, i.e. the dissimilarity between the expression patterns of those two markers. In practice, we use L1-norm as the dissimilarity measure in this work. Assume we have a training set *S* and a validation set *P*. Initially, we set the loadings as follows,

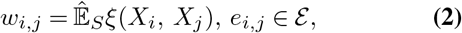

where 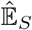 denotes the empirical expectation over *S*. After initializing, 𝒱 we assign a *cost* to each node in by a non-negative function *q* : 𝒱 → ℝ, where *q*(*v*) denotes the cost of node *v*, ∀*v* ∈ 𝒱. Also, we distinguish edges by if they are activated, an activated edge by means it associates at least one node whose associated RV is in 𝒳_obs_ and goes out from the node in 𝒳_obs_. When *e*_*i,j*_ is activated, we denote *κ*_*i,j*_ = 1. That is, *κ*_*i,j*_ = 1 if *X*_*i*_ ∈ 𝒳_obs_ and 0 otherwise. For a node whose associated RV in 𝒳_pred_, the node cost is assigned by the minimal edge loading of activated edges it associates with, and 0 for a node whose associated RV is in *𝒳*_obs_. That is, ∀*X*_*i*_ ∈ 𝒳

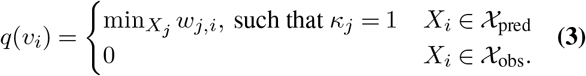

We start with 𝒳_in_ containing only one RV, whose associated node had the minimal node cost. Then, in each iteration, the predictor *f*^*t*^ is trained by ERM and predictions are made for RVs in 𝒳_pred_. Our goal is to have node costs measuring the unpredictability of their associated RVs, and edge loadings measuring the risk occurring when predicting the value of one connected node from the other observed connected node, i.e. the dissimilarity between predicted and real patterns. We update the loadings by the predictor’s generalization performance on a held-out validation set *P* after training a new predictor, i.e. ∀*X*_*i*_, *X*_*j*_ ∈ 𝒳, *X*_*i*_ ≠ *X*_*j*_,

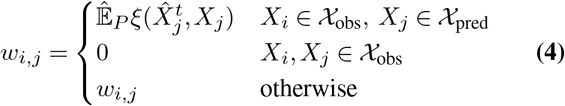

where 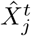 denotes the prediction of *X*_*j*_ from predictor *f*_*t*_. Note that the loading of an edge from a predicted node to an observed node will not be updated since the unpredictability is only regarding the direction from observation to prediction. In general, the initial dissimilarity measure between the expression patterns of a pair of markers is an approximate upper bound of the unpredictability (i.e., the value that would result if the prediction is trivially made by outputting the input patterns). The algorithm aims to gradually reduce the overall unpredictability among all markers of interest by iteratively including informative markers into 𝒳_obs_, as illustrated in Algorithm 2.

#### Algorithm 1 Predictive Marker Identification

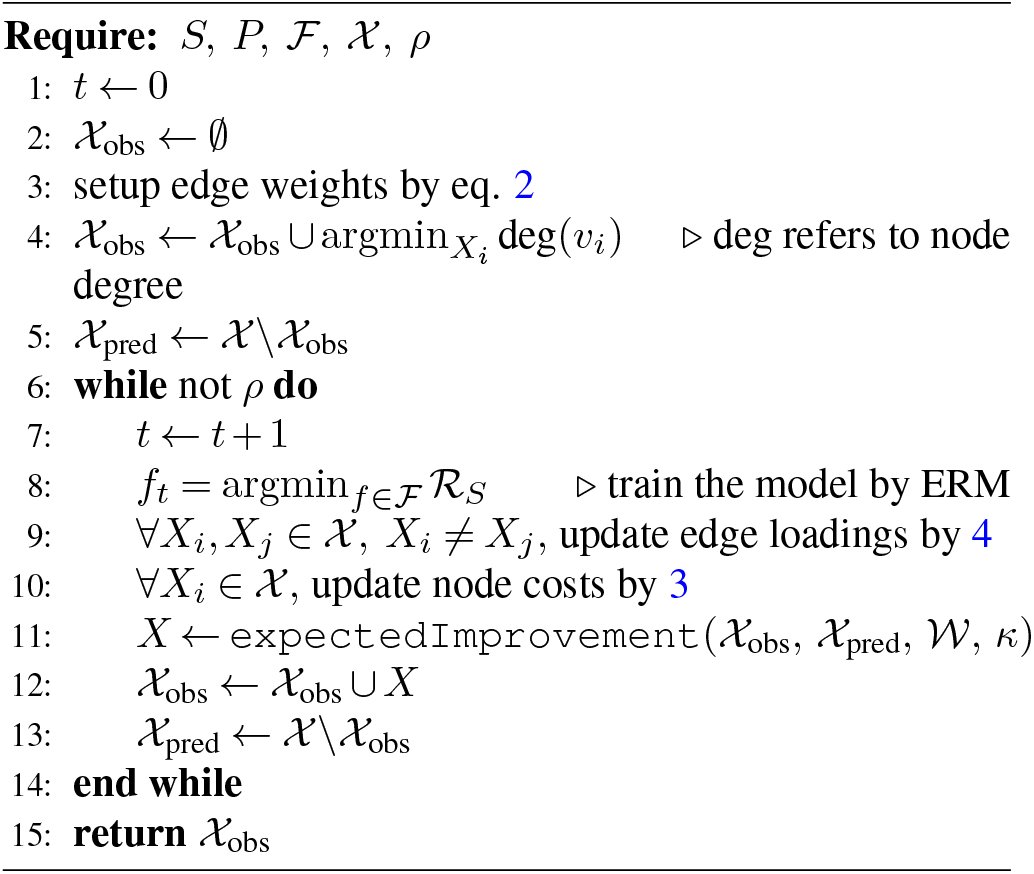

In practice, we selected the RV that would most decrease the summation of the node costs Σ_*i* ∈ [|*𝒳*|]_ *q*(*v*_*i*_). After selecting a new marker into the predictive set, we retrain the predictor using the updated set of selected markers. We illustrated the full algorithm in Algorithm 1.

#### Algorithm 2 subroutine: expectedImprovement

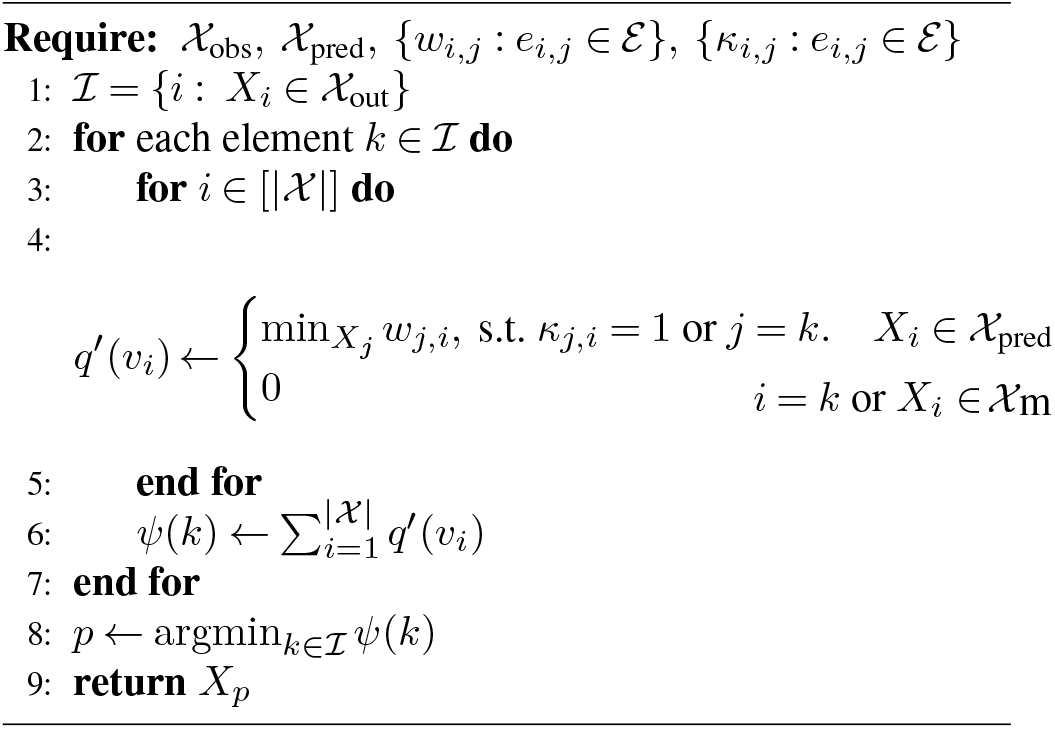

### Multipanel

The single panel setting assumes the training and validation sets consist of images with all protein markers of interest observed. However, this is not always true when the number of markers of interest is large. For example, suppose we have 200 markers of interest, but current imaging technology only allows us to have up to 60 markers in a single image. Then, the single-panel setting will fail as it requires the samples in the *S* and *P* to contain all RVs in 𝒳 in order to initialize 𝒢. We, therefore, extended the algorithm to solve this problem.

For simplicity, we first consider the case that the markers of interest, 𝒳, can be measured using only two panels. We denote the markers in the two panels by 𝒳_*A*_ and 𝒳_*B*_ respectively, where 𝒳_*A*_ ∪ 𝒳_*B*_ =𝒳. Here, we assume the sizes of both panels are less than or equal to the maximum number of markers that can be imaged simultaneously. Also, we expect there will be a set of *overlapping markers* appearing in both panels, namely 𝒳_*M*_ = 𝒳_*A*_ ∩ 𝒳_*B*_ *=* ∅. For markers in every single panel, we can first set up individual sub-graphs using the same approach as for the single panel in the single-panel setting described above. We then consider completing the whole graph by inferring the loadings of edges between RVs in different panels. Recall that the edge loadings are initialized by the dissimilarity of two RVs and updated by their unpredictability from one to the other RV associated with their connecting nodes. If we assume the unpredictability is a valid norm (e.g. L1-norm), for any two markers not in the same panel, we can get an upper bound on the edge loading between *X*_*i*_ and *X*_*j*_ by the triangle inequality, *w*_*ij*_ ≤*w*_*iq*_ + *w*_*qj*_, for *X*_*q*_ ∈ 𝒳_*M*_. By doing so, the distributed version degenerates to the single-panel setting for which we already had an approximate solution above. Each round, two predictors are retrained with respect to markers in 𝒳_*A*_ and 𝒳_*B*_ respectively using the training sets *S*_*A*_ and *S*_*B*_, depending on the panel where the last marker was selected. In the multipanel setting, the cost of edge *e*_*i,j*_ is initialized by the following three cases: when both *X*_*i*_ and *X*_*j*_ both in the overlap set 𝒳_*M*_, the edge loading is the average dissimilarity (measured by *ξ*) between two associated RVs among two panels, as 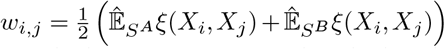 ; when two nodes are both in a single panel and not both in the overlap set, we have 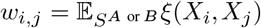; and lastly, when two nodes locate in separated panels, we apply the triangle inequality as 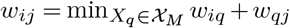

It is then trivial to extend the method to a multipanel setting. That is, consider the whole set of markers 𝒳 being partitioned into *m* subsets associated with *m* panels, i.e. 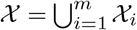 and for each subset, there exists at least one other subset, their *i* = *k* or *X*_*i*_ intersection is not empty (as in the two-panel case they share some overlap markers), which indicates the whole graph of marker RVs has no disconnected compartments. By doing so, for every two markers not in the same panel, we can find at least one path in the graph connecting their associated RVs, and therefore we can complete the whole set of edge loadings by chaining the triangle inequality and find the minimal loading value if there exist multiple paths. In practice, finding paths is realized by standard depth-first search. If a marker is predicted by multiple predictors, the predictions will be averaged.

### Data preparation and machine learning

To illustrate the single-panel setting, we used spleen and lymph node CODEX image datasets from data published by the HuBMAP project (14). These contained 8 and 9 multichannel images of different tissue regions respectively. We split the datasets into training, validation, and test sets of 4:2:2 for the spleen dataset and 4:2:3 for the lymph node dataset. Each image in those data sets contains 29 measured biomarkers.

We used HuBMAP datasets for large and small intestine image datasets containing 16 different multichannel images, where each image contains 46 measured biomarkers, to test the multipanel settings. For both datasets, 4 images were held out as a separate test set. The remaining 12 images, were evenly split into training and validation sets.

The intensities of each channel were normalized for each CODEX image using the same normalization method as (13) for spleen and lymph node datasets, as follows

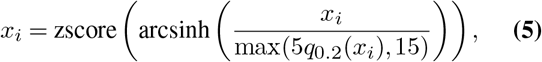

where *x*_*i*_ denotes the i-th channel of the image, *q*_0.2_ denotes the 20-th percentile, and zscore 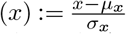. For small and large intestine datasets, we directly applied zscore normalization, since the signal in these two datasets is relatively weak and sparse. Since the size of each CODEX image was very large, during training time, patches (channel number ×192 ×192) were randomly cropped from a CODEX image as the input or target of the predictor. However, during testing, the patches were cropped as a sliding window (not randomly) from a single CODEX image, and the original size image was recovered by stitching those patches.

We used convolutional neural networks as predictors in this work. In particular, the network architecture was a U-Net (16) with skip connections (17) which has been widely used in modern computer vision applications. The predictor was trained to minimize the empirical mean square error (MSE) using an Adam optimizer with a learning rate of 10^−4^. The validation set was used to monitor the training process and the predictor with the lowest validation loss was selected.

As the predictive set selection required re-training predictors each round (Line 8 in Algorithm 1), the predictor in round *t* + 1, *f*_*t*+1_, inherited the trained weights from round *t, f*_*t*_, for *t >* 0. Since the predictor architecture of *f*_*t*+1_ had an additional input channel and one fewer output channel compared to *f*_*t*_, the weights of the selected channel in the input layer were randomly initialized in *f*_*t*+1_ and the weights of this channel in the output layer of *f*_*t*_ were not inherited. We randomly initialized the weights of *f*_1_ following a normal distribution.

### Quality measures

Our first assessment is the overall un-predictability of, 𝒳i.e. the sum of node costs, measured by the L1-norm. A second is the predictor’s performance on the test set in terms of reconstruction MSE and Pearson Correlation Coefficient (PCC) between synthetic and real images. We also examined the quality of single-cell level predictions. We first segmented individual cells from test images. For spleen and lymph node datasets, we directly used segmented masks from HuBMAP. For intestine datasets, we segmented the cells using DeepCell Mesmer (18). We then create matrices (for both real and synthetic images) whose rows refer to individual cells, columns refer to the markers, and each entry refers to the average intensity of a marker for a particular cell. We also created another single-cell profile referred as the correlation profile, whose entries are PCC between a pair of marker patterns within a cell region. That is, the columns of this matrix refer to every pair of markers of interest. We measured the difference between the matrices resulting from real and synthetic images using the normalized Frobenius norm.

### Feature vectors of immunohistochemical image patches

For an even smaller size of each subpanel, we have to partition the markers of interest into several subpanels. We designed the sub-panels according to the similarity of the patterns of markers. Since predictors will be trained for each sub-panel respectively, similarities within a panel gives the best chance of their being predictable from each other. The similarity was measured by feature vectors resulting from a neural network classifier trained on immunohistochemical image patches. There are 334 and 336 whole-side images of 41 markers available for colon and small intestine in the HPA (for markers without HPA immunohistochemical images, we randomly added them into sub-panels). Here, for each whole slide immunohistochemical image, we decomposed it into 2 channels referring to the protein marker and tissue background using the algorithm described in (19), and randomly cropped it into patches from regions with high expression. For each tissue, we randomly split all patches into training and validation sets, and trained a ResNet-50 based image model to learn to classify protein expression patterns according to their associated protein markers. We then collected all feature vectors associated with each image patch (from both training and validation sets) from the second-to-last layer of the classifier and performed PCA to reduce their dimensionality to 2. Next, we grouped the protein markers according to the centroids of their feature vectors in the PC coordinates, resulting in 5 subpanels for both tissues.

