## Supplementary Information for "Expanding the coverage of spatial proteomics"

### Expanding the coverage of spatial proteomics: Supplementary Information

September 8, 2023

#### 1 Datasets

##### Spleen

###### Training

1. HBM495.VWBD.428
2. HBM825.KFFT.669
3. HBM396.BXSQ.568
4. HBM863.FDNH.844

###### Validation

1. HBM455.PWQW.883
2. HBM967.TGDD.996

###### Test

1. HBM455.XDQS.993
2. HBM594.PWXG.764

##### Lymph Node

###### Training

1. HBM622.JXWQ.554
2. HBM992.RHJW.288
3. HBM268.NKXB.243
4. HBM557.SGTC.262

###### Validation

1. HBM938.TNNT.879
2. HBM522.BSZT.385

###### Test

1. HBM347.PSLC.425
2. HBM997.PVCF.629
3. HBM685.TBGN.663

##### Large Intestine

###### Training

1. HBM334.QWFV.953
2. HBM353.NZVQ.793
3. HBM424.STVV.842
4. HBM462.JKCN.863
5. HBM622.STKS.394
6. HBM575.THQM.284

###### Validation

1. HBM683.NRPR.962
2. HBM729.XTBN.693
3. HBM739.HCWP.359
4. HBM429.LLRT.546
5. HBM438.JXJW.249
6. HBM439.WJDV.974

###### Test

1. HBM742.NHHQ.357
2. HBM792.FFJT.499
3. HBM938.KMNW.825
4. HBM964.FPNH.767

###### New batch of experiments

###### Training

1. HBM396.FNQW.543
2. HBM423.MMGW.744
3. HBM352.MDZF.598
4. HBM292.FCMS.497
5. HBM953.LMWQ.235
6. HBM725.QFKT.594

###### Validation

1. HBM685.PCCJ.427
2. HBM753.VDXD.934
3. HBM634.MSKL.575
4. HBM994.KDNT.678

###### Test

1. HBM946.NWTV.278
2. HBM524.VWGB.378
3. HBM886.NTZN.682
4. HBM494.VNTQ.422

##### Small Intestine

###### Training

1. HBM443.LGZK.435
2. HBM466.XSKL.867
3. HBM666.RBCG.529
4. HBM284.SBPR.357
5. HBM676.QVGZ.455
6. HBM785.FJVT.469

**Validation**

1. HBM687.SJLD.889
2. HBM934.KLGL.584
3. HBM893.MCGS.487
4. HBM845.VMSZ.536
5. HBM899.KTQM.246
6. HBM945.FSHR.864

**Test**

1. HBM394.VSKR.883
2. HBM727.DMKG.675
3. HBM953.KMTG.758
4. HBM996.MDQH.988

**New batch of experiments**

**Training**

1. HBM443.XPDK.549
2. HBM795.GWKV.825
3. HBM475.DXDC.532
4. HBM644.QKZS.857
5. HBM334.RPTP.997
6. HBM233.GTZN.466

**Validation**

1. HBM423.QJJR.545
2. HBM398.SWKV.256
3. HBM837.RHNC.533

**Test**

1. HBM587.VTDD.789
2. HBM829.QZQP.626
3. HBM735.PSTF.274
4. HBM962.BFPH.344

### 2 Selected Markers

#### Spleen

CD5, DAPI, SMActin, CD31, CD15, Vimentin, CD20

#### Lymph node

FoxP3, CD3e, PanCK, CD163, HLA-DR, Vimentin, CD8, CD20, CD1c, LYVE-1, CD11c, CD45RO

#### Large intestine

##### Single panel

CD16, CD49a, Hoechst, aSMA, Vimentin, CD90, CD57, SOX9, ITLN1, CDX2, CD44, MUC1, CD127, Cytokeratin, CD45RO, CD34, CD19, HLADR, CD4

##### Multipanel

CD127, CD21, CD138, Hoechst, CD49a, CD163, Vimentin, CD90, aSMA, SOX9, CD57, MUC1, CD15, CDX2, CD44, HLADR, CD7, CD34, CD3

#### Small intestine

##### Single panel

CD68, CD45RO, CD4, SOX9, aSMA, Ki67, CD3, CD90, Cytokeratin, CD44, CD49a, MUC2, CD49f, Hoechst, CD31, CD38, ITLN1, CD45, CD16

##### Multipanel

SOX9, CD45RO, CD49a, CD4, aSMA, CD68, DRAQ5, CD117, CDX2, CD45, MUC2, ITLN1, CD21, Hoechst, CD66, Vimentin, CD163, CD44, CD31

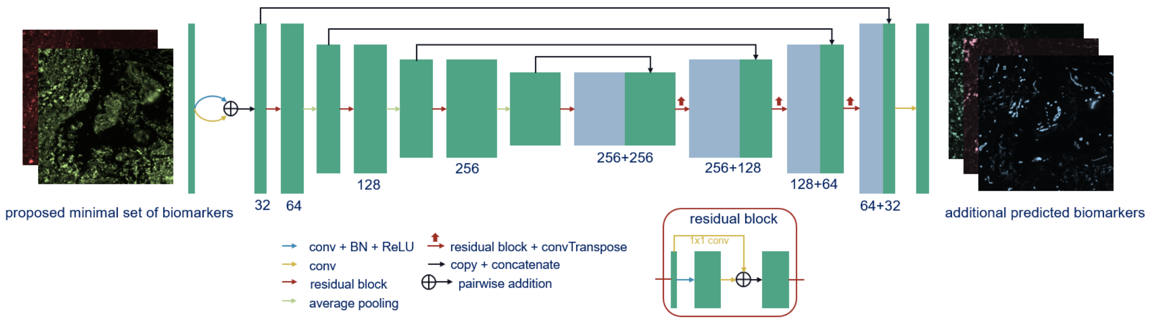


Figure S1: Architecture of the convolutional neural network used as the predictor in this paper.

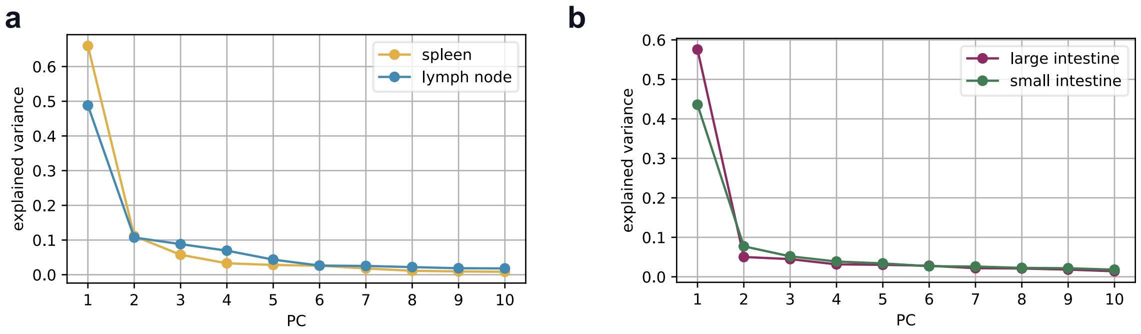


Figure S2: Variance explained by each principal component (PC) for single-cell level expression of spleen and lymph node cells (**a**) and large and small intestine cells (**b**) is shown for the test sets of the four tissues.

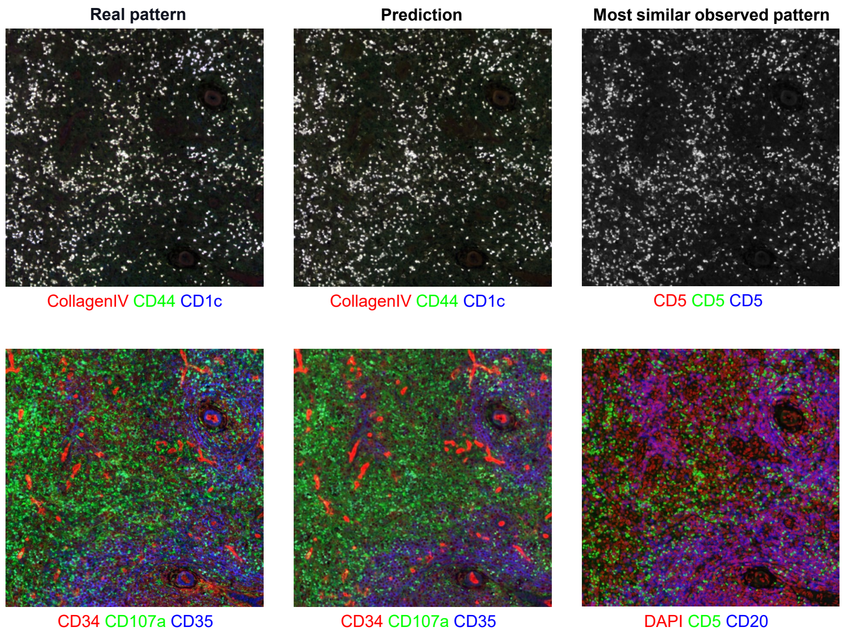


Figure S3: Expression patterns of markers in the selected predictive set that are most similar to the corresponding image patches shown in the first two columns. For example, for the lymph node, CD5 is the marker selected whose pattern is most similar to CollagenIV, CD44, and CD1c's. These illustrate that accurate prediction can be made even for markers with different patterns than the input patterns. The image patches are shown from the spleen test set.

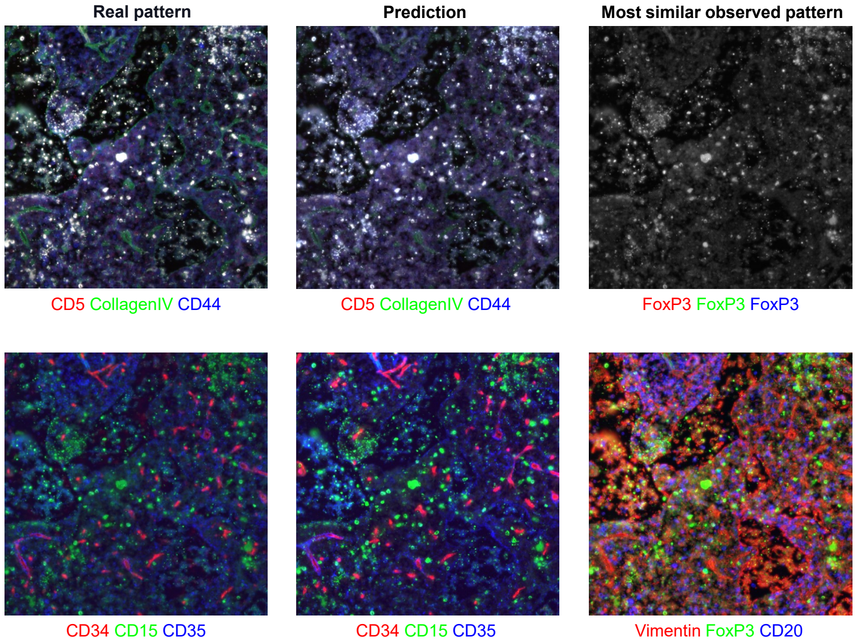


Figure S4: Similar to Figure S3, the most similar input expression patterns to the patterns shown in the first two columns. The image patches are shown from the lymph node test set.

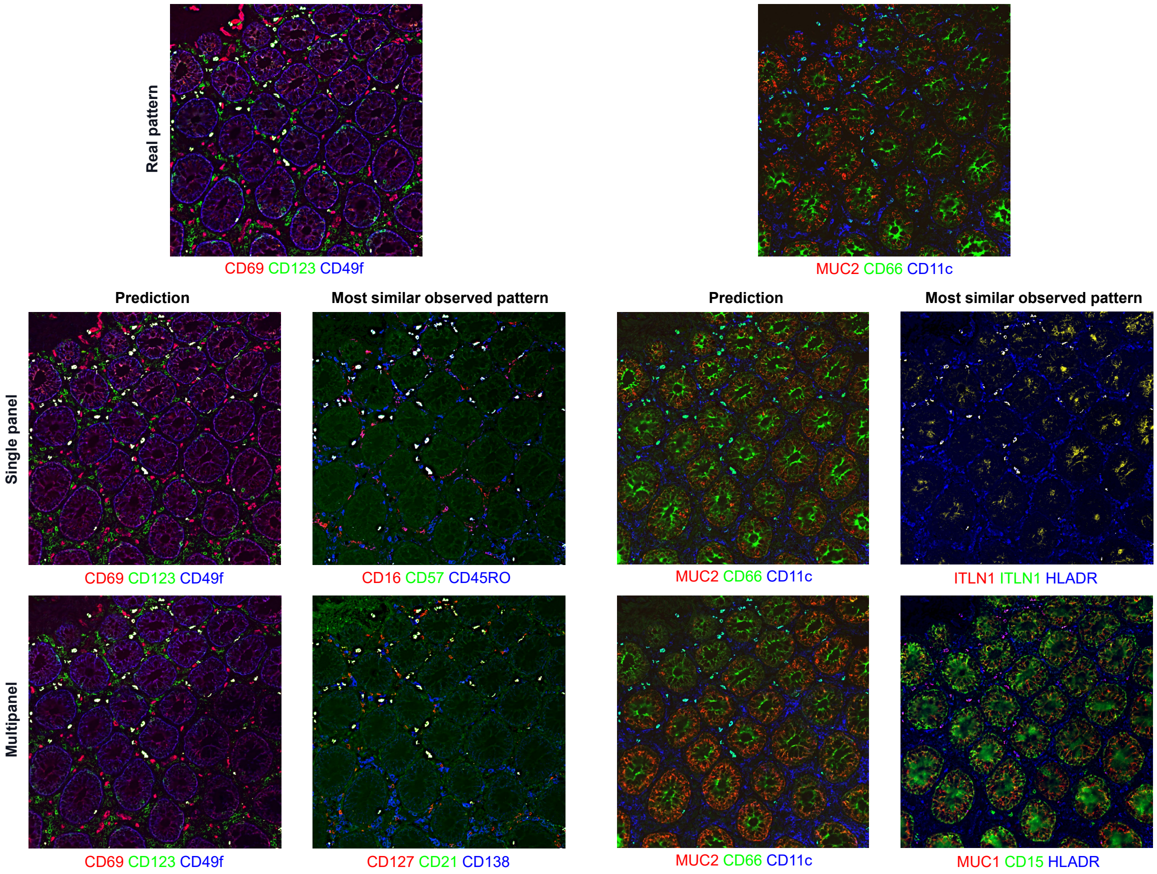


Figure S5: Similar to Figure S3, the most similar input expression patterns to the corresponding real and predicted patterns. The image patches are shown from the large intestine test set.

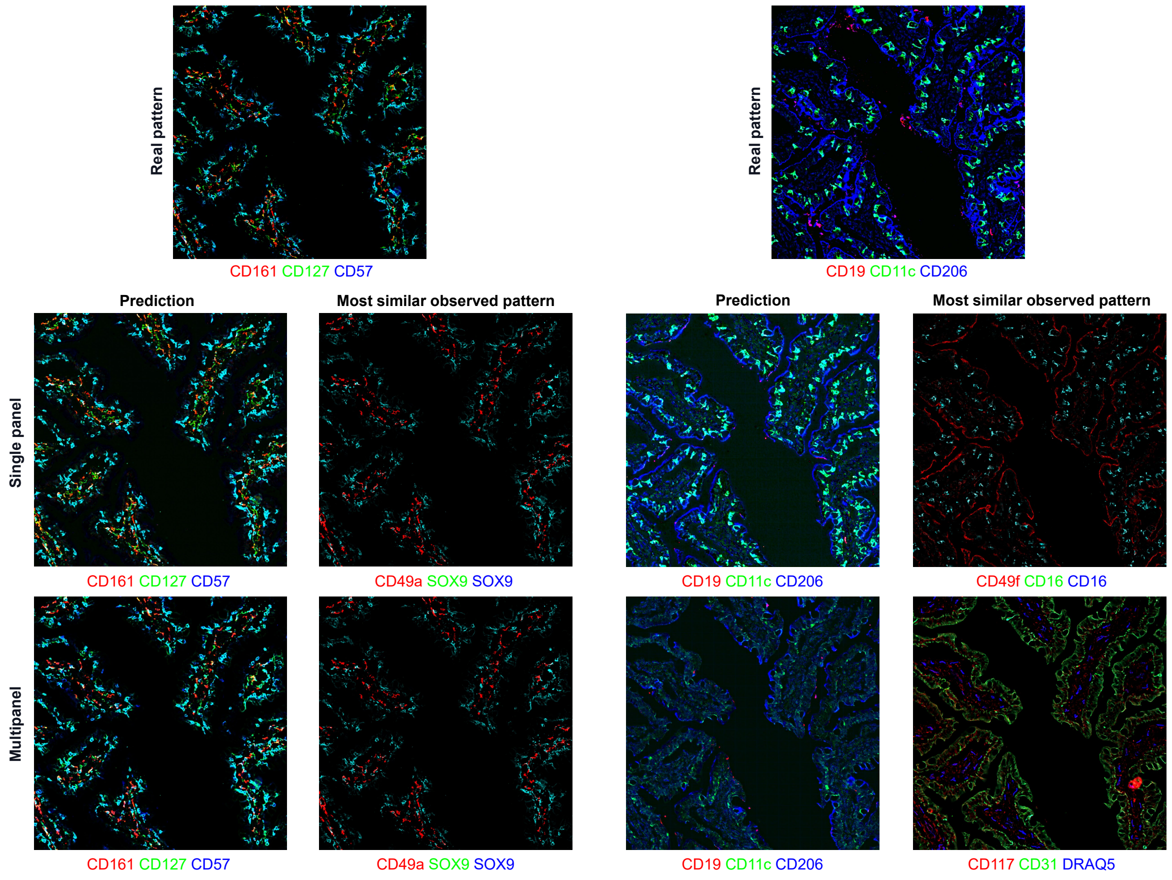


Figure S6: Similar to Figure S3, the most similar input expression patterns to the corresponding real and predicted patterns. The image patches are shown from the small intestine test set.

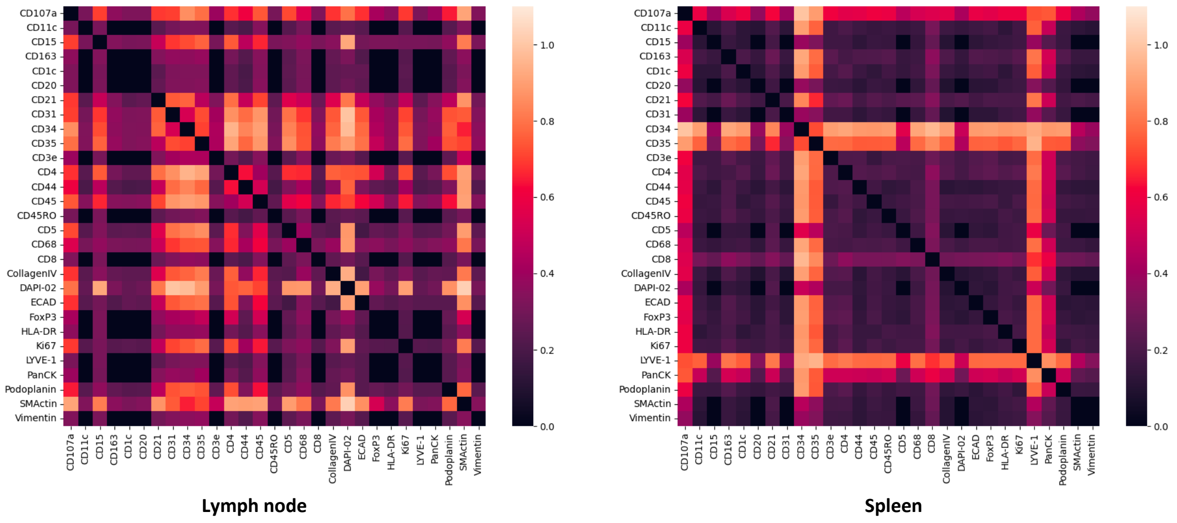


Figure S7: Heatmap showing the final graph from lymph node and spleen dataset using single panel method. Each matrix represents the final edge weights of each model after the selection process stops. Each matrix entry indicates the final weight of the edge connecting a pair of biomarkers at the corresponding coordinates. Although Figure 1 in the main text represents the graph edge bidirectionally, here we consider both directions with the same weight for the sake of simplicity, which means edges in both directions are updated simultaneously in the selection process.

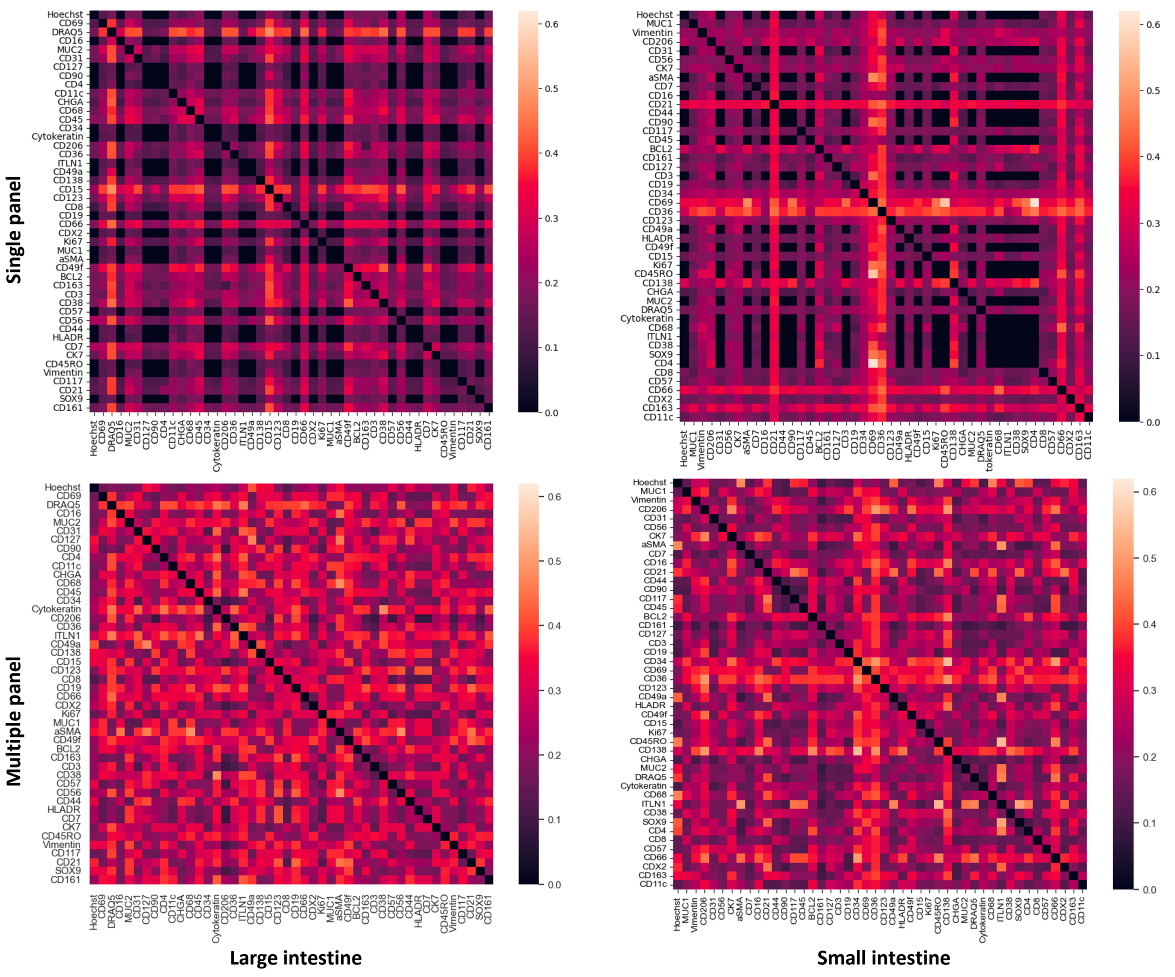


Figure S8: Similar to Figure S7, heatmap showing the final graph from the large intestine and small intestine dataset using single panel method and multi-panel method.
